# Switch in parasitic and autotrophic-dominated protist assemblages coupled to seasonal oligotrophic-mesotrophic gradients in the sunlit layer of a subtropical marine ecosystem

**DOI:** 10.1101/2024.10.16.618647

**Authors:** Yoav Avrahami, Raffaele Siano, Max Rubin-Blum, Gil Koplovitz, Nicolas Henry, Colomban de Vargas, Miguel J. Frada

## Abstract

Protists are major functional players in the oceans. Time-resolved epipelagic protist successions remain poorly described in subtropical ecosystems, limiting current understanding of food web dynamics and responses to environmental changes in these major world-ocean regions. Here, we used amplicon sequencing data and their trait-based annotation to examine the seasonality of planktonic protists in the subtropical Gulf of Aqaba (northern Red Sea). Temperature and nutrients were identified as major drivers of succession. Marked seasonal shifts in protists were detected. The warm, stratified oligotrophic period spanning through spring and summer were dominated by heterotrophs, including parasitic Syndiniales. By contrast, nutrient influx during deep convective-mixing in winter triggered a progressive shift to photoautotrophic communities dominated by few chlorophyte genera. Ephemeral phytoplankton blooms were detected during the mixing-to-stratification transition. Deeper winter mixing resulted in larger blooms dominated by diatoms and coccolithophores relative to chlorophytes prevalent during shallower mixing. This illustrates the impact of mixing-depth in the development and compostion of blooms. Comparisons with oceanwide rDNA datasets indicate that the oligotrophic protist assemblages from the Gulf of Aqaba resemble those from warm, open-oceans. This work provides a detailed assessment of the seasonality in protist communities and dominant functional strategies in a coastal subtropical planktonic ecosystem.

## Introduction

Planktonic protists represent the majority of eukaryotic diversity in the sunlit layer of the ocean, comprising an extensive and still largely unknown variety of functional roles that are determinant to food web structuring and global biogeochemical processes (Falkowski et al., 2004; de Vargas et al., 2015; Lima-Mendez et al., 2015; Worden et al., 2015; Guidi et al., 2016). Auto- and mixo-trophic protists (eukaryotic phytoplankton) are main constituents of marine primary production (Behrenfeld et al., 2001), while heterotrophic protists serve as important grazers of bacteria and phytoplankton and are consumed by higher trophic levels (Sherr & Sherr, 2002). Other marine protists establish symbiotic associations with other microbes and larger organisms, along a continuum from mutualism to parasitism, and can act as a major regulator of plankton population dynamics and community succession (Coats, 1999; Anderson & Harvey, 2020; Bass et al., 2021; Blindheim et al., 2023). Much of the protistan diversity has not yet been characterized or cultured, and field studies of protist assemblages complemented by physiological lab experiments are rare (Caron et al., 2009; Keeling, 2013; Keeling & Campo, 2017), although critical for a better understanding of the functional basis of marine ecosystems. High-throughput amplicon sequencing of biodiversity marker genes (barcodes) can provide a detailed account of the taxonomic diversity of a given sample, while offering new insights into functional diversity and the complexity of food web structure in marine ecosystems (de Vargas et al., 2015; Lima-Mendez et al., 2015; Massana et al., 2015; Pernice et al., 2016; Ramond et al., 2019; Meyneng et al., 2024).

Subtropical, oligotrophic marine ecosystems >40% of the world’s ocean, and account for about 25% of the marine primary production (Longhurst, 1995; Polovina et al., 2008; Duarte et al., 2013). However, current descriptions of the succession patterns of planktonic protist communities in these ecosystems are limited. Most studies in oligotrophic, subtropical regions are based on oceanographic transects covering large spatial but limited time scales (de Vargas et al., 2015; Lima-Mendez et al., 2015; Malviya et al., 2016). Fewer studies have provided comprehensive time and taxonomic resolutions to protist diversity patterns in these regions (Pasulka et al., 2013; Blanco-Bercial et al., 2022). Moreover, subtropical oceans are predicted to warm up and expand pole-wards due to global warming, which exerts thermal stress on local species at the limits of thermal tolerance (Chust et al., 2014; Cheng et al., 2022) and drive the replacements of high-latitude communities. Therefore, better descriptions of community turnover along seasonal time scales are necessary to improve our understanding of the dynamics of taxonomic and functional diversity in subtropical ecosystems, as well as to generate baseline information to detect future ecosystem shifts in response to climate change. In this context, we aimed to examine the diversity and community succession of planktonic protists in the Gulf of Aqaba (GoA) in the northern Red Sea as a model subtropical ecosystem. The GoA is a marginal, warm and narrow marine basin surrounded by deserts that reaches a depth of >700m, a few kilometers from the surrounding shorelines (Kienast & Torfstein, 2022). However, during about half of the year (April-November) the GoA resembles an open-ocean gyre ecosystem from the perspective of epipelagic physical, chemical and biological profiles (Berman & Gildor, 2022). The water column is markedly stratified (with a shallow, typically <30 m wind-driven mixed layer) and oligotrophic. Cyanobacteria numerically dominate the phytoplankton communties and deep-chlorophyll maximum forms around the depth of 100m on the verge of the nutricline (Lindell & Post, 1995; Zarubin et al., 2017). Peak surface temperatures and the most pronounced thermocline gradients are detected in August (Zarubin et al., 2017). By contrast, during fall-winter (November-March), the system shifts to turbulent, mesotrophic conditions, as a consequence of intense deep-convective mixing. This mixing progresses as a result of surface cooling and reaches hundreds of meters in depth (c.a. 200m to >700m), homogeneizing water column and markedly enhancing nutrient fluxes to the photic layer as an inverse function of temperature. These deep-mixing events are enabled by a relatively narrow temperature gradient across the water column in the Red Sea (Carlson et al., 2014). The GoA deep mixing dynamics sets the stage for spring blooms developing over a short period (<1 month) at the onset of the stratified period. The magnitudes of these blooms are unusual for subtropical, oligotrophic seas and comprise diverse diatom assemblages (Zarubin et al., 2017; Avrahami et al., 2024). They decay rapidly likely due to nutrient limitation as stratification intensifies (Zarubin et al., 2017; Berman & Gildor, 2022). As such, the GoA provides a good opportunity to examine with high temporal resolution the succession patterns of planktonic protist communities inhabiting subtropical marine systems, in response to environmental gradients and interannual variations in mixing depth.

We used rDNA amplicon sequencing (18S rDNA, V4-region, using eukaryotic universal primers (Stoeck et al., 2010)) to generate datasets of amplicon sequence variants (ASVs), that were then annotated taxonomically, together with phenotypic and functional traits associated with taxa (Ramond et al., 2019). The processed dataset was then used to determine the succession patterns of both protist taxonomic and functional diversity as a key component to ecosystem dynamics (Weisse et al., 2016; Ramond et al., 2019; Singer et al., 2021). Finally, we integrated our ASV dataset into comparable data collected across the world’s surface ocean (Pesant et al., 2015), aiming to test whether similar protistan communities are observed in other parts of the world ocean, supporting the use of the GoA as a model open-ocean subtropical ecosystem. Overall, our study provides a detailed synthesis of how both temporal and spatial environmental gradients developing seasonally and interannually influence protist community dynamics in a subtropical marine ecosystem.

## Experimental Procedures

### Study site and sampling

Between November 2020 and May 2022, we collected near-surface water samples from a depth of ∼2m at the open ocean site station A (29°28’N, 34°55’E) in the northern GoA (bottom depth ∼700 m). During mixing (November-March), samples were collected twice a month. At the peak of mixing and during the spring bloom (March-April), samples were collected weekly or twice per week to capture the rapid phytoplankton bloom dynamics. At each time point, seawater was transferred into acid-washed carboys using a manual pump installed in a boat and pre-filtered by 500 µm mesh to avoid large zooplankton. In addition, a Sea-Bird SBE 19 CTD (conductivity, temperature, depth, Sea-Bird Scientific) was deployed to obtain the physical parameters of the water column. In the stratified summer (late April-August) samples were collected both from the surface and the deep chlorophyll maximum (DCM) using 12 L Niskin bottles (General Oceanics) installed on the Sea-Bird carousel and coupled with Sea-Bird SBE 19+ CTD (Sea-Bird Scientific). Mixed layer depth (MLD) was calculated according to (Zarubin et al., 2017), as the depth in which there is a difference of 0.2 °C of potential temperature, compared to 3m depth. Further analysis was processed shortly upon return to the lab (30-60 min).

### DNA extraction and sequencing

In the lab, duplicates of ∼4.4 L seawater were filtered onto 0.2 µm Sterivex filters (Merck Millipore), flash-frozen in liquid N_2_ and stored at -80 °C until DNA extraction. DNA was extracted using the NucleoSpin Plant II kit (Macherey-Nagel), after the protocol used by the Tara Oceans project (Alberti et al., 2017). The extraction began with incubation of the filter with proteinase K and lysozyme (120 min at 56 °C). Lysate was transferred to a clean tube and was treated according to the kit’s guidelines. Blank samples of clean filters were used as negative controls. Following the extraction, DNA quality and concentrations were measured with QFX fluorometer (Denovix), using the dsDNA quantification assay (high sensitivity). After DNA extraction and quality test, samples were shipped to the Genomics and Microbiome Core facility at Rush University for PCR and sequencing by MiSeq v2 chemistry (Illumina). Primers targeted the V4 domain of 18S-rDNA (∼380 bp), as used by the TARA Ocean project (TAReuk. Forward: 5’-CCAGCASCYGCGGTAATTCC-3’, Reverse: 5’-ACTTTCGTTCTTGATYRA-3’) (Stoeck et al., 2010). Sequence data are available at GenBank (accession PRJNA1169359).

### Sequence data processing and analyses

Sequences were processed with Qiime 2 (Bolyen et al., 2019). Demultiplexed paired-end reads were trimmed with Cutadapt (Martin, 2011) without indels, to remove primer sequences, followed by quality control and denoising by DADA2 (Callahan et al., 2016). Sequences were assigned to ASVs and annotated by the protist reference database PR2 (version 4.14) (Guillou et al., 2013) using BLAST by a cutoff of 90% (Camacho et al., 2009). 7,370 unique ASV’s were identified after the removal of unassigned reads and annotations of the phylum Metazoa. Duplicates per each sampling point were averaged and yielded 43 sampling points which included 1,180,541 reads in total. Saturation of taxa diversity of over 99.9% in all sampling points was verified by rarefaction curves with the R package vegan (R Core Team (2021); Oksanen et al., 2022) (Supp. Fig. 1a).

**Figure 1.**
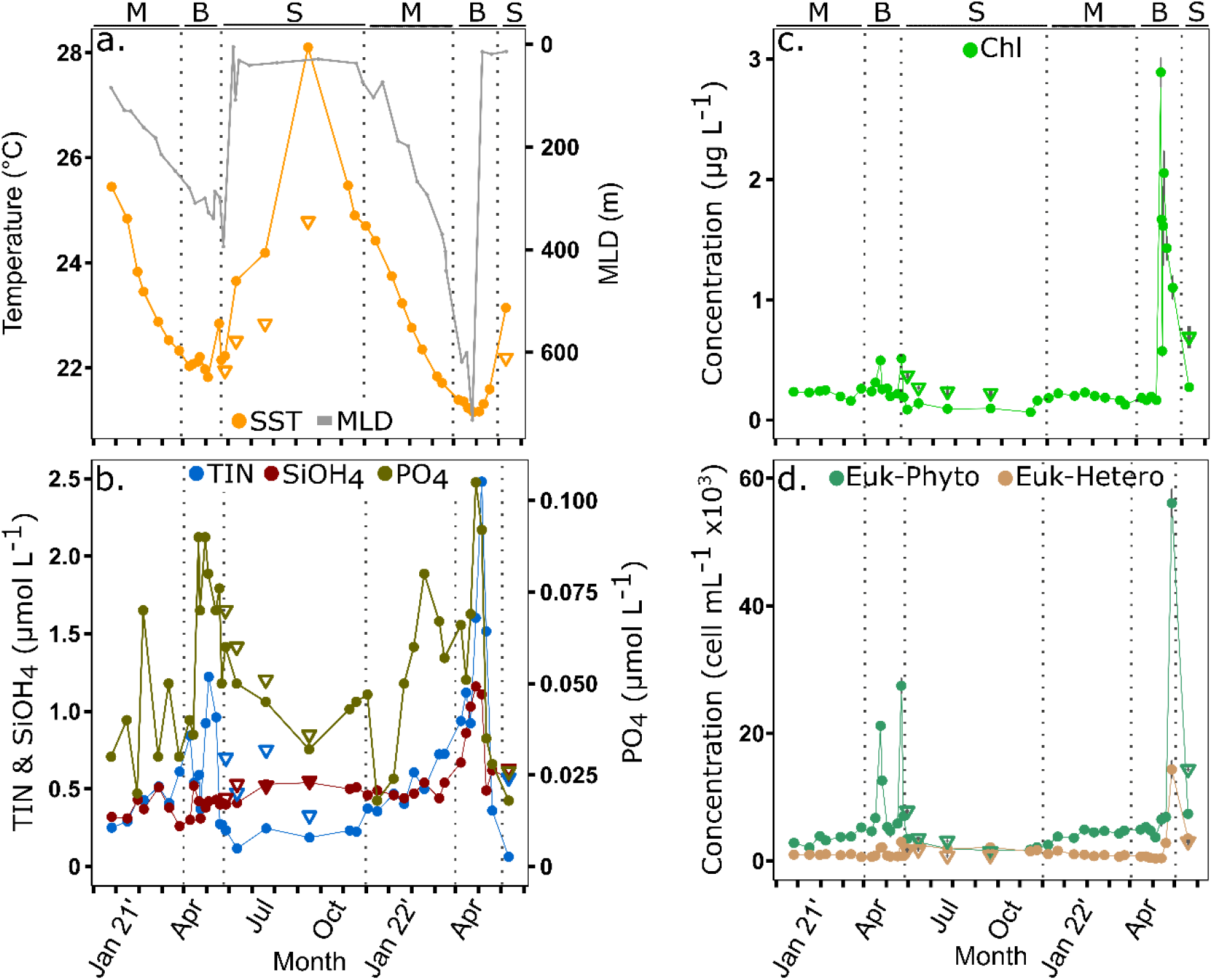
Physical, chemical and biological variables in the Gulf of Aqaba. Measurements at the Deep chlorophyll maximum (DCM) during the stratified season are represented as triangles.(a) Temperature at the Sea surface (SST) and DCM, and mixed layer depth (MLD). (b) Macronutrients: Total Inorganic Nitrogen (TIN), Silica, and Phosphate. (c) Chlorophyll a. (d) Total eukaryotic phytoplankton (Euk-Phyto) and eukaryotic heterotrophs (Euk-Hetero) as determined by flow cytometry. Grey bars in C and D represent the standard deviation (n=3, n=2, respectively). The seasons are separated by dashed lines and noted by letters in the top of each panel: M=mixing, B=bloom, S=summer stratification.

Richness was calculated based on the ACE index which distinguishes rare and abundant species in its formulation with the R package phyloseq (McMurdie & Holmes, 2013). Turnover (temporal beta diversity index) was calculated with the R package codyn (Hallett et al., 2016) as species appearance and disappearance divided by total species between two time points. This analysis was performed on one time point only for each month, to avoid bias of different time intervals between sampling points. Community composition explained by environmental drivers was based on relative abundance values and carried by Redundancy Analysis (RDA) with the vegan package in R (Oksanen et al., 2022).

To compare the GoA and Tara Oceans datasets, we used the “Tara Oceans (2009-2013) rDNA 18S V4 ASV table” (Delage et al., 2023), extracting only samples comprising full size-fraction (>0.8 µm), and from mid or low latitudinal regions were included, which limited comparisons with samples from the North and South Atlantic and Pacific and one sample from the mid-Red Sea collected during winter.

Trait data curated by (Ramond et al., 2019) encompassed morphology, physiology, life-cycle and trophic mode of protists, and covered 2,006 taxa with 32 traits. We selected the 18 most informative traits and manually assigned taxa in our study on the genus level, as protistan functional traits are overall conserved within genera (Finlay, 2004; Litchman et al., 2007; Violle et al., 2011). When there was a conflict within a genus for values of a trait, it was assigned as NA. Functional assignment in the GoA applied for 243 genera (average 80% of the total genera in the GoA). Then, based on Gower distance method (Gower, 1971), we clustered genera with similar trait composition into functional groups. The number of groups was determined based on K-means clustering and Principal Coordinate Analysis (PCoA) which placed the selected genera based on their trait similarity on an ordination space and highlighted 8 distinct groups. Next, we characterized each group based on the main shared traits and defined the groups according to size and trophic mode, and to a lesser extent by cell cover. The obtained groups were: pico/nano-auto, silicified-auto, micro-auto (mixotrophs), pico-parasite, pico-hetero (including different heterotrophs as phagotrophs, parasites, saprotrophs, and osmotrophs), pico-phago, micro-phago, and armored-micro-phago. Functional richness index (FRic) measures the volume of a convex hull formed by the presence of traits in an ordination space (Villéger et al., 2008) and was calculated by using the R package mFD (Magneville et al., 2022). We compared functional richness to genera richness, hence the same data of 243 selected genera was used for this analysis (and not total ASV’s as used to measure taxonomic richness). To form the functional composition of the GoA with the global Tara Ocean data, we assigned additional genera that were highly represented in Tara Ocean stations (over 70% of total reads per station) to obtain a total of 432 genera from both data sets. Following the same clustering protocol, 6 groups emerged similar to the GoA-only functional groups, whereas the smaller number of groups stemmed from the merging of several size classes into groups of a wider size range. The outcome groups were: pico-nano-micro-auto, silicified-auto, pico-parasite, pico-phago, nano-micro-phago, and armored-micro-phago.

### Nutrient analyses

Total inorganic nitrogen (TIN, NO_2_+NO_3_+NH_4_), silicate (SiO_2_), and orthophosphate (PO_4_) were determined. For the analysis of NO_2_, NO_3_, silicate, and phosphate, sub-samples of 10 mL were transferred in triplicates to 15 mL Falcon tubes and analyzed with Flow Injection autoanalyzer (Quik-Chem 8500, LACHAT Instruments). Every run was calibrated by using commercial standards with at least 5 sub-standards to cover the whole working range. For ammonium (NH_4_) measurements, quadruplicates of 4 mL were transferred to 15 mL Falcon tubes and analyzed fluorometrically (Hoefer).

### Chlorophyll a

Triplicates of 300 mL seawater were filtered onto Whatman glass fiber filters (GF/F, 25 mm diameter, 0.7 µm pore size). The filters were incubated in 90% acetone (Carlo Erba Reagents) buffered with saturated MgCO3 (Sigma-Aldrich), over-night (4 °C in darkness), and analyzed fluorometrically for chlorophyll a (Trilogy, Turner Designs) (Jeffrey and Humphrey, 1975).

### Flow cytometry

Triplicates of 4 mL seawater were collected onto Falcon tubes. Samples were preserved with 0.25% Glutaraldehyde and stored at 4 °C for 30 minutes. Then, samples were flash-frozen in liquid N_2_ and kept at -80 °C until analysis. Analysis was performed by Attune Nxt flow cytometer (Thermo Fisher Scientific), and targeted phytoplankton groups of eukaryotes and cyanobacteria (Marie et al., 2014).

## Results

### Physical, chemical and biological variations across seasons

Marked seasonal changes in environmental parameters were detected between November 2020 and May 2022 (Fig. 1a-c). These data served as a basis to identify 3 distinct seasonal periods: 1) the stratified period (late April-November) characterized by progressively higher sea surface temperatures, marked water-column stratification and low nitrate (0.19 µmol L^-1^) and chlorophyll levels (<0.14 µg L^-1^). Warming was also detected at the DCM, where nutrients and chlorophyll were higher than at the surface; 2) the winter-mixing period (November-March) was characterized by progressively declining sea-surface temperature and concomitant water-column convective mixing. As a consequence, nitrate (0.47 µmol L^-1^), phosphate and chlorophyll (0.21 µg L^-1^) increased relative to summer; 3) the bloom period (April-May) where annual maxima for nutrients and chlorophyll were detected. The bloom coinceded with the onset of stratification, following the deepest mixing depth. Maximal mixing depth varied annually, In 2021 mixing reached about 300m deep, while in 2022 it surpassed 700m deep. As a consequence the concentrations of nitrate (1.3 µmol L^-1^), silicate (0.85 µmol L^-1^) and chlorophyll (1.1 µg L^-1^) in April 2022 were much higher than those measured in April 2021 (0.67 µmol L^-1^ nitrate, 0.4 µmol L^-1^ silicate, and 0.3 µg L^-1^ chlorophyll).

Variations in the concentration of total eukaryotic phytoplankton (Euk-Phyto) as detected by flow cytometry closely matched the chlorophyll profiles (Fig. 1d). During the stratified period at the surface the concentrations of Euk-Phyto were about 3,000 cells mL^-1^. The concentrations increased to about 3,870 cells mL^-1^ with the onset of winter-mixing and peaked during the blooms. Higher cell counts were detected in the 2022 bloom (>12,000 cells mL^-1^) relative to 2021 (>10,000 cells mL^-1^). Total eukaryotic heterotrophs concentrations were considerably lower than photoautotrophs at all times (Fig. 1d). A decrease in concentration was detected through the winter-mixing as compared to summer surface, with cell concentrations averaging 966 and 2,177 cells mL^-1^, respectively. The Peak of heterotroph concentrations coincided with the bloom of 2022 (2,840 cells mL^-1^), whereas the bloom in 2021 contained relatively low heterotrophs (1,307 cells mL^-1^).

### Seasonal variations in protist diversity

We observed marked seasonal variations in the main planktonic protist Phyla (Fig. 2). Overall Dinoflagellata and Chlorophyta were the most predominant Phyla, interchanging in prevalence between seasons Dinoflagellata was more common during the stratified period, accounting for up to 65% of the total ASV reads. They declined during the mixing and bloom periods to about 30% of total ASVs, but rapidly recovered with the onset of stratification. Of the Dinoflagellata ASV read abundance, ∼36% were assigned to undefined Dinophyceae (Dinophyceae_XXX), ∼25% to Syndiniales Dino-groups I and II clade, and ∼10% to a group of dinoflagellate genera composed of *Gyrodinium, Prorocentrum* and *Tripos* (Supp. Fig. 2a). About 24% of Dinoflagellata sequences comprised a variety of taxa with taxonomic annotations at various levels. By contrast, Chlorophyta were rare at the surface during the stratified season (< 5% of the ASVs), but gradually increased through the mixing period to ∼25-50% read abundance during the bloom (Fig. 2). *Ostreococcus, Micromonas* and *Bathycoccus* were the most common Chlorophyta genera. *Ostreococcus* represented about 85% of Chlorophyta ASV read abundance during the mid-mixing and bloom periods but was much rarer on the surface during the stratified period and early mixing (<5% of Chlorophyta ASVs). *Micromonas* and *Bathycoccus* dominated the Chlorophyta ASV read abundance at the surface during the stratified seasons (Supp. Fig. 2b). At the DCM during the stratified period, the relative contribution of Chlorophyta was higher than the surface, and *Ostreococcus* contributed 15-80% of Chlorophyta ASV read abundance. Noticeably, the relative contribution of Chlorophyta was lower common during the bloom of 2022, coinciding with an increase in Ochrophyta and Haptophyta.

**Figure 2.**
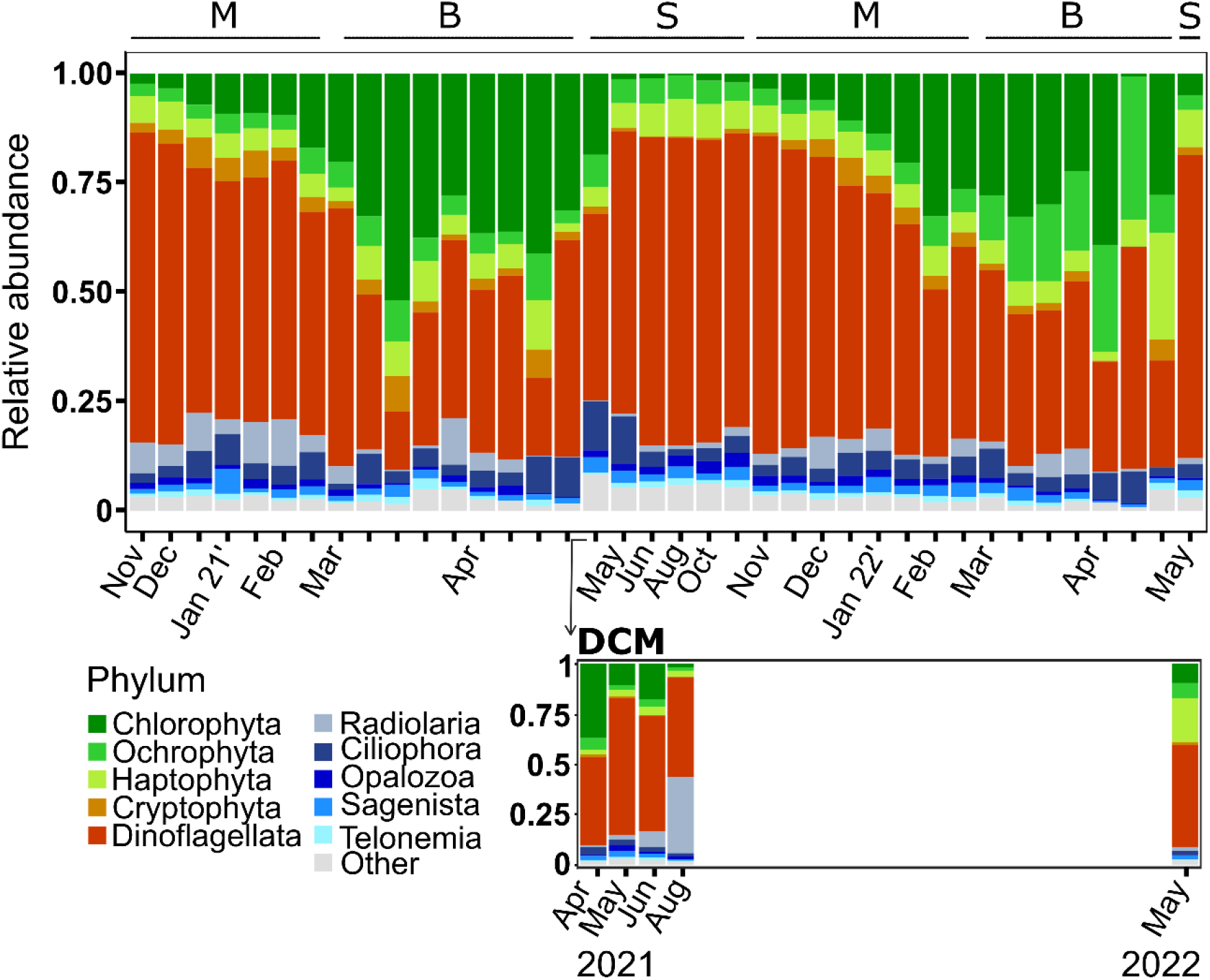
Main Protist Phyla in the Gulf of Aqaba. Averaged (duplicates) relative abundance of the top 10 most abundant Phyla in surface waters (upper panel) and the corresponding main phyla at the DCM during summer (lower panel), based on ASV’s average reads (duplicates). Letters in the top panel represent seasonal periods: M=mixing, B=bloom, S=summer stratification.

Ochrophyta and Haptophyta represented about 13% of ASVs across seasons. Peak of relative abundances were distinct during the blooms (Fig. 2). *Aureococcus* and diatoms (*Skeletonema, Thalassiosira* and species from the Family *Mediophyceae*) were the most common Ochrophyta during the later mixing period and bloom. *Pelagomonas* prevailed during the early mixing and DCM. Representatives of the Marine Ochrophyta group (MOCH-2 (Massana et al., 2014)) and a large fraction of unannotated sequences (>65%) prevailed in surface waters during the stratified period and the early mixing (Supp. Fig. 2c). The genera *Chrysochromulina* and the coccolithophore *Gephyrocapsa* were the most common Haptophyta, the latter specifically during the 2022 bloom. Finally, Cryptophyta represented in general <5% of the total ASVs during the mixing and bloom season, and were rarer during the stratified season. Heterotrophic protists represented 10-25% of ASV sequences across all time points. Radiolaria, Ciliophora and Opalozoa were the most represented Phyla. Radiolaria were more prevalent during winter (except for a peak in August at the DCM). Ciliophora peaked during the bloom and the early stratified period (Fig. 2, Supp. Fig. 2d,e).

### Protist alpha diversity patterns

Total protist richness (ACE Index) and diversity (Shannon Index, H’) displayed covarying seasonal patterns (Fig. 3a). Higher richness and diversity were detected during the stratified season (ACE ∼700; H’ >5). A gradual decline was detected during the mixing period. Minimal levels were detected during the bloom (ACE <200; H’ <4). A rapid recovery in richness and diversity was detected at the onset of the stratified period. Richness patterns for the main Phyla detected (Dinoflagellata, Chlorophyta, Ochrophyta and Haptophyta) followed similar seasonal oscillation. Richness was typically higher during the stratified season at the surface (Fig. 3b). However, the relation between ASV sequence abundance and Richness was dissimilar between Phyla. Read abundance and richness closely covaried for Dinoflagellates. This was not the case for the other groups. Namely in Chlorophyta the highest sequence abundance during late-mixing and bloom seasons coincided with the lowest Richness values (Fig. 3b). Co-variation patterns in Ochrophyta and Haptophyta were less clear (Fig. 3b). However, peak of Richness for both groups were detected during the stratified season and the 2022 bloom.

**Figure 3.**
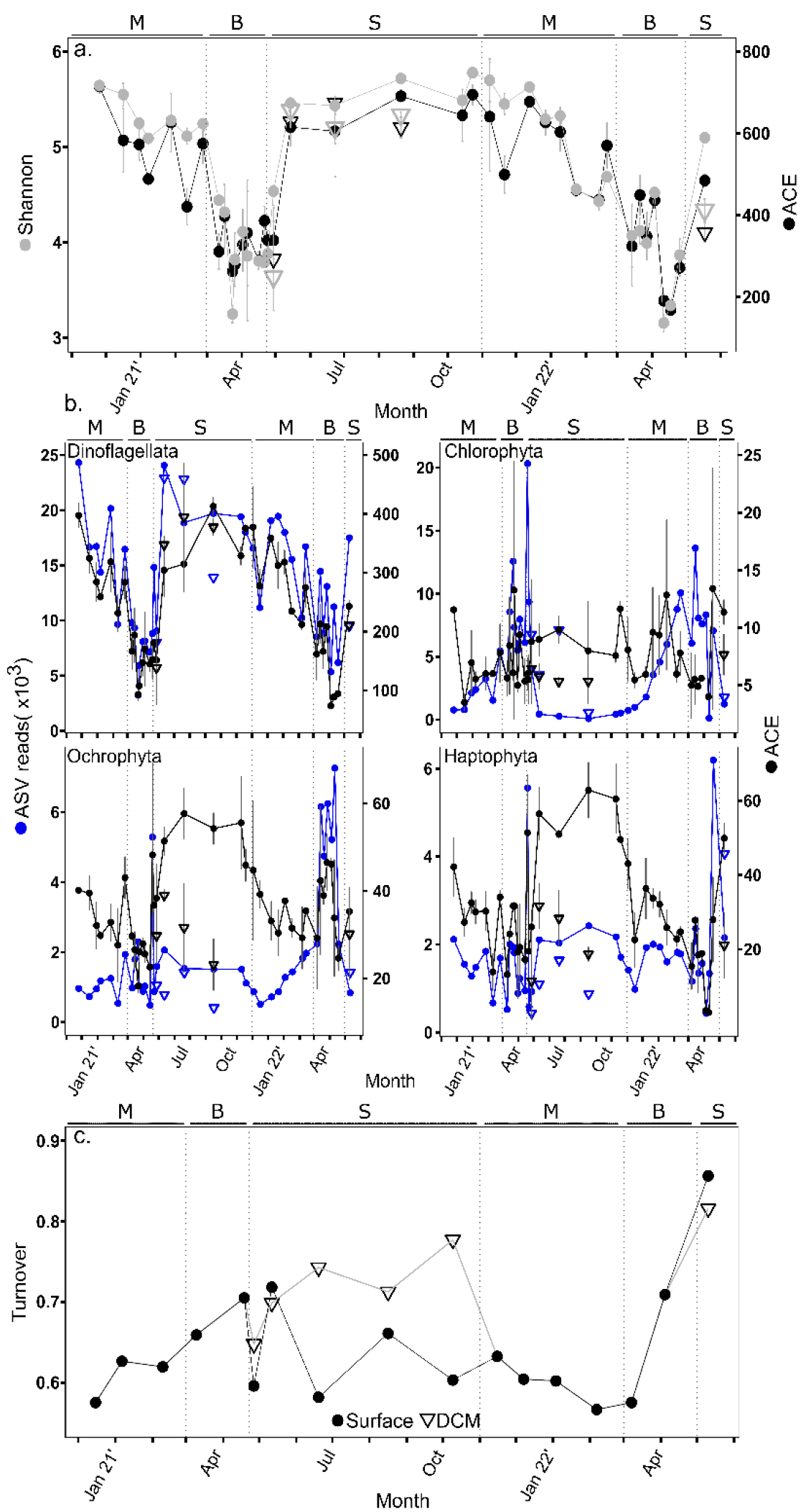
Protist diversity indices. (a) Diversity (Shannon index) and richness (ACE index) of total ASV reads. Grey bars represent standard deviation (n=2). (b) Total ASV reads (blue) and richness (ACE index, black) for the main autotrophic phyla. Grey bars represent standard deviation (n=2). (c) Temporal turnover index. This index detects differences between time points in appearance and disappearance of ASV’s. For (a), (b), and (c), circles represent samples from sub-surface waters (2m depth), while triangles represent samples from the deep chlorophyll maximum (DCM). The seasons are separated by dashed lines and noted by letters in the top of each panel: M=mixing, B=bloom, S=summer stratification.

The turnover index was used as an estimator of the rate of ASV replacement over time (Fig. 3c). Higher turnover rates were detected during the progression of the bloom period, notably in 2022, and particularly at the DCM during the stratified season (0.81 on average). Lower turnover values were detected in surface waters during the stratified season (0.77 on average) and the mixing season (0.72 on average).

### Relationships between protist communities and environmental variables

Redundancy Analysis (RDA) of protist assemblages (Fig. 4) showed a compact cluster from the stratified seasons and early mixing seasons associated with higher temperature, and lower nutrient levels. By contrast, samples from the later mixing stages and particularly the bloom associated with low temperature and higher nutrients were more dispersed, indicating higher variations in assemblages. Samples collected at the DCM during the stratified seasons frequently overlapped with winter and bloom samples. The most distinct protist assemblages were detected during the 2022 bloom period.

**Figure 4.**
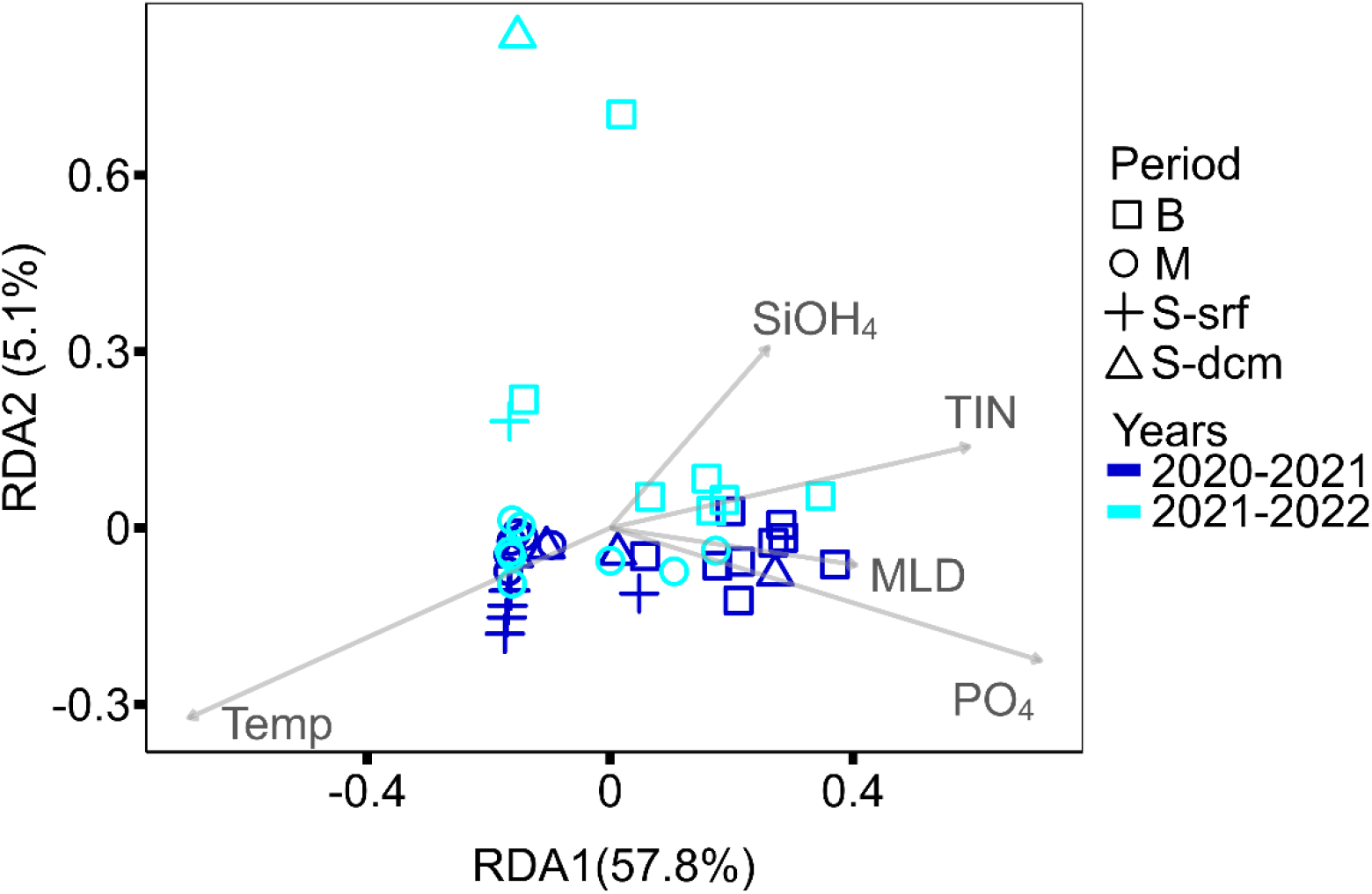
Redundancy Analysis (RDA) ordination based on the relative abundance of protistan communities in the Gulf of Aqaba. Symbols represent the seasonal phase of each sample: B= bloom, M= mixing, S-srf= summer surface, S-dcm= summer dcm. Colors represent the year of sampling: blue= 2020-2021, cyan= 2021-2022. Grey arrows represent significant environmental variables (p <0.05): Temp= temperature, SiOH4= silicic acid, TIN= total inorganic nitrogen, MLD= mixed layer depth, PO4= phosphate. In parenthesis next to each axis label is the percentage of explained variability by the axis.

### Seasonal variation in functional groups of protists

Analyses of the functional traits of protist ASVs with clear taxonomic identification enabled the distinction of eight functional clusters (Fig. 5; Supp. Fig. 3): 1) pico/nano-autotrophs largely composed of small chlorophytes and other small phytoplankton groups, 2) silicified-autotrophs composed of diatoms; 3) micro-autotrophs; 4) pico-parasites composed of Syndiniales and other Dinophyceae; 5) pico-heterotrophs corresponding to a variety of functional types including osmotrophic, parasitic, saprotrophic organisms; plus two groups of heterotrophic genera with predatorial lifestyles that could be differentiated base on size: 6) pico-phagotrophs composed by heterotrophic Marine Stramenopiles (MAST); 7) micro-phagotrophs containing a variety of ciliates and heterotrophic dinoflagellates. A final group 8) was identified as armored-micro-phagotrophs, composed of Radiolaria (Supp. Table 1).

**Figure 5.**
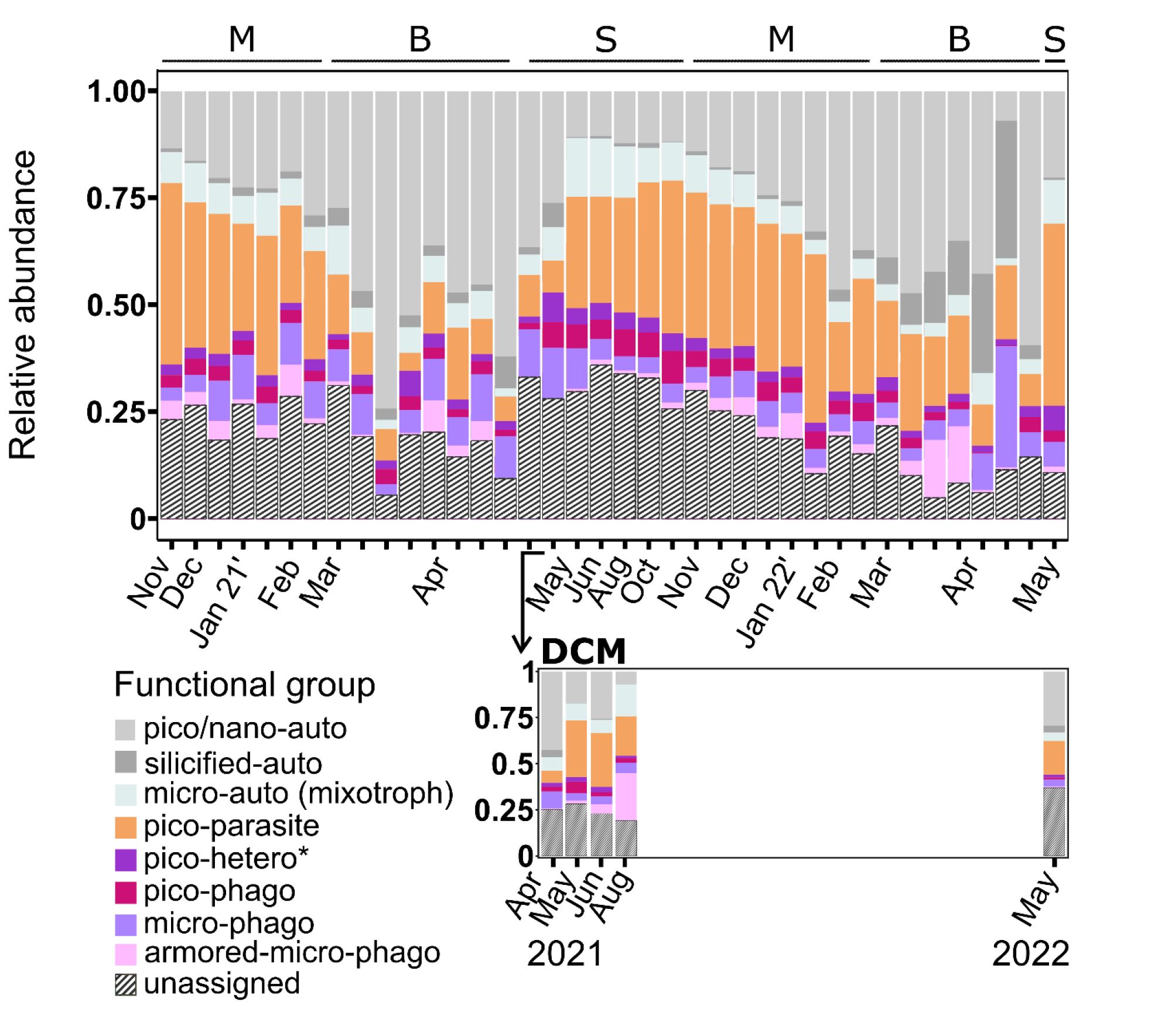
Composition of protistan functional groups over seasonal gradients in the Gulf of Aqaba. Grey-shaded bars represent autotrophic groups, and colored bars represent heterotrophic groups. Averaged (duplicates) relative abundance of functional groups at surface waters (upper panel) and the corresponding functional groups at the DCM during summer (lower panel), based on 243 most abundant genera in the GoA. Letters in the top panel represent seasonal periods: M=mixing, B=bloom, S=summer stratification.

Noticeably, an important fraction of the diversity remained unassigned to any functional category. These represented 5-20% of the diversity during the bloom, and 20-30% during the stratified and mixing seasons.

Overall, pico/nano-autotrophs and pico-parasites were the most prevalent functional groups identified (Fig. 5), broadly matching the seasonality of parasitic dinoflagellates (Syndiniales and other parasitic taxa) and Chlorophyta, respectively (Supp. Fig. 2). The first was more common during the stratified season and the second was in the mixing and bloom. Micro-autotrophs (many of which likely display mixotrophic attributes) were also more common during the stratified period. Silicified autotrophs were more common during the bloom, especially in 2022 where they reached 36% of the total read abundance (compared to 8% during the bloom of 2021). Pico-heterotrophs, pico-phagotrophs, micro-phagotrophs and armored-micro-phagotrophs inter-varied in density over seasons, but as a unit represented typically 15-20% of the functional diversity in the Golf of Aqaba.

The FRic index was used to test the relation between genetic and functional richness and to assess for each sampling point the volume of functional space occupied by genera (Villéger et al., 2008). A positive correlation between functional richness and genetic richness was detected (R^2^=0.61, Fig. 6). Time points from the mixing and stratified seasons were characterized by the highest genetic and functional diversity, while by contrast, bloom time points had overall lower values, although a wider richness range.

**Figure 6.**
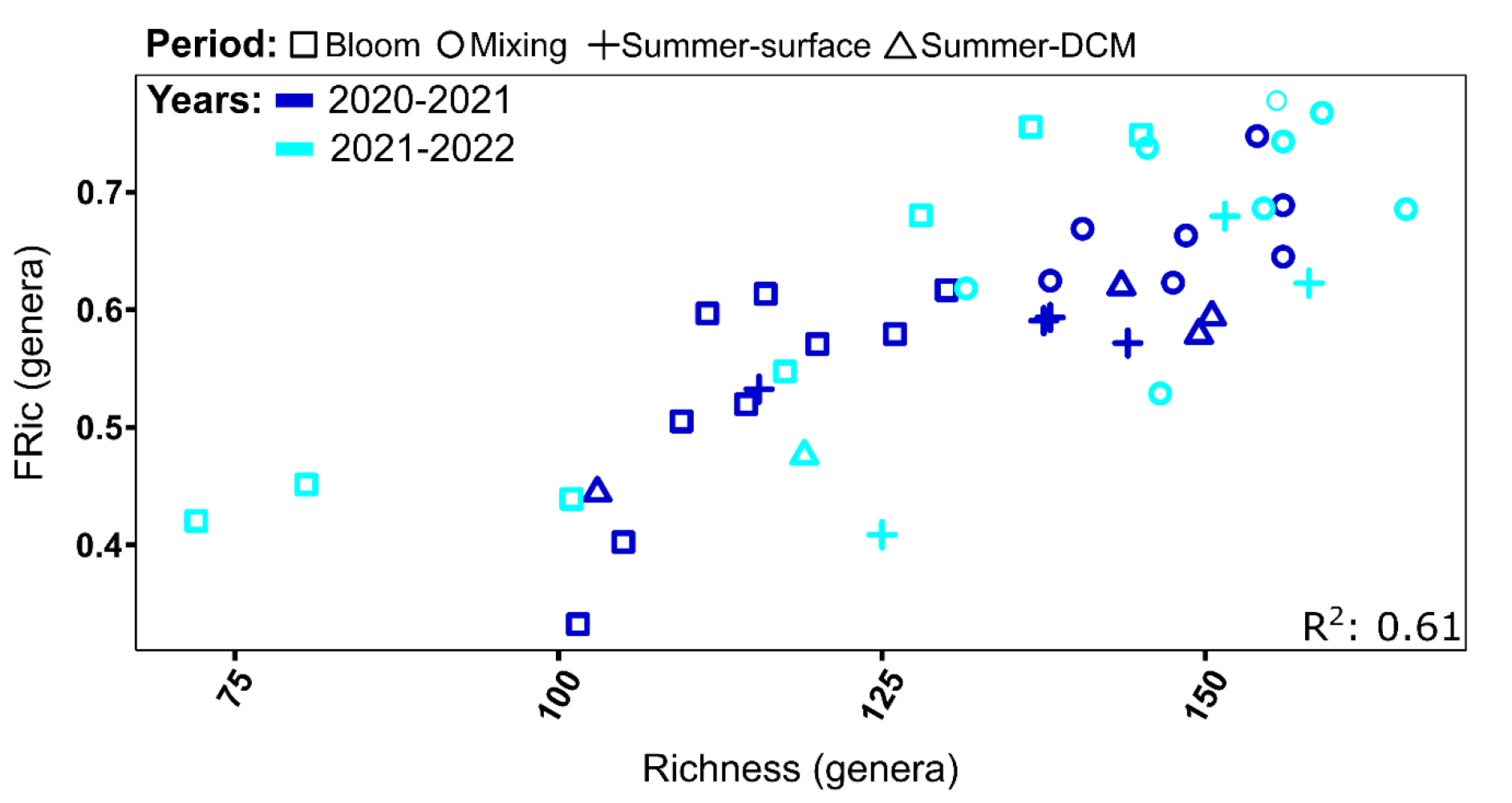
Patterns of taxonomic versus functional richness. Each sampling point was measured for taxonomic richness (x-axis) and functional richness (Fric, y-axis), based on an analysis performed at the genus taxonomic level and including the 243 top protist genera in the GoA. Shapes represent different seasonal periods: Square=bloom, Circle=mixing, Plus=summer-surface, Triangle=summer-DCM. Colors represent sampling years: blue= 2020-2021, cyan=2021-2022. R squared was calculated by a linear model of the 2 variables.

### Comparison of protist assemblages between the GoA and other (sub)topical regions from the world ocean

We integrated our data from the GoA into the 18S-V4 amplicon dataset from the *Tara* Oceans circumglobal expedition (Pesant et al., 2015). Higher variability was observed in our seasonal samples from the GoA, which spread along the RDA1 axis (Fig. 7b). Among the GoA variability, samples collected in surface waters during the stratified season, some from the DCM as well as early mixing periods markedly associated with samples collected globally in warm, oligotrophic subsurface and DCM waters from the North Atlantic and North and South Pacific oceans. By contrast, samples from the GoA collected during the late mixing and bloom periods, with rare exceptions, did not cluster with any of the Tara Oceans samples from the world’s open ocean. (Fig. 7b). This could be mainly explained by a large prevalence of Haptophytes from the genus *Chrysochromulina* in the GoA (Supp. Fig. 4).

**Figure 7.**
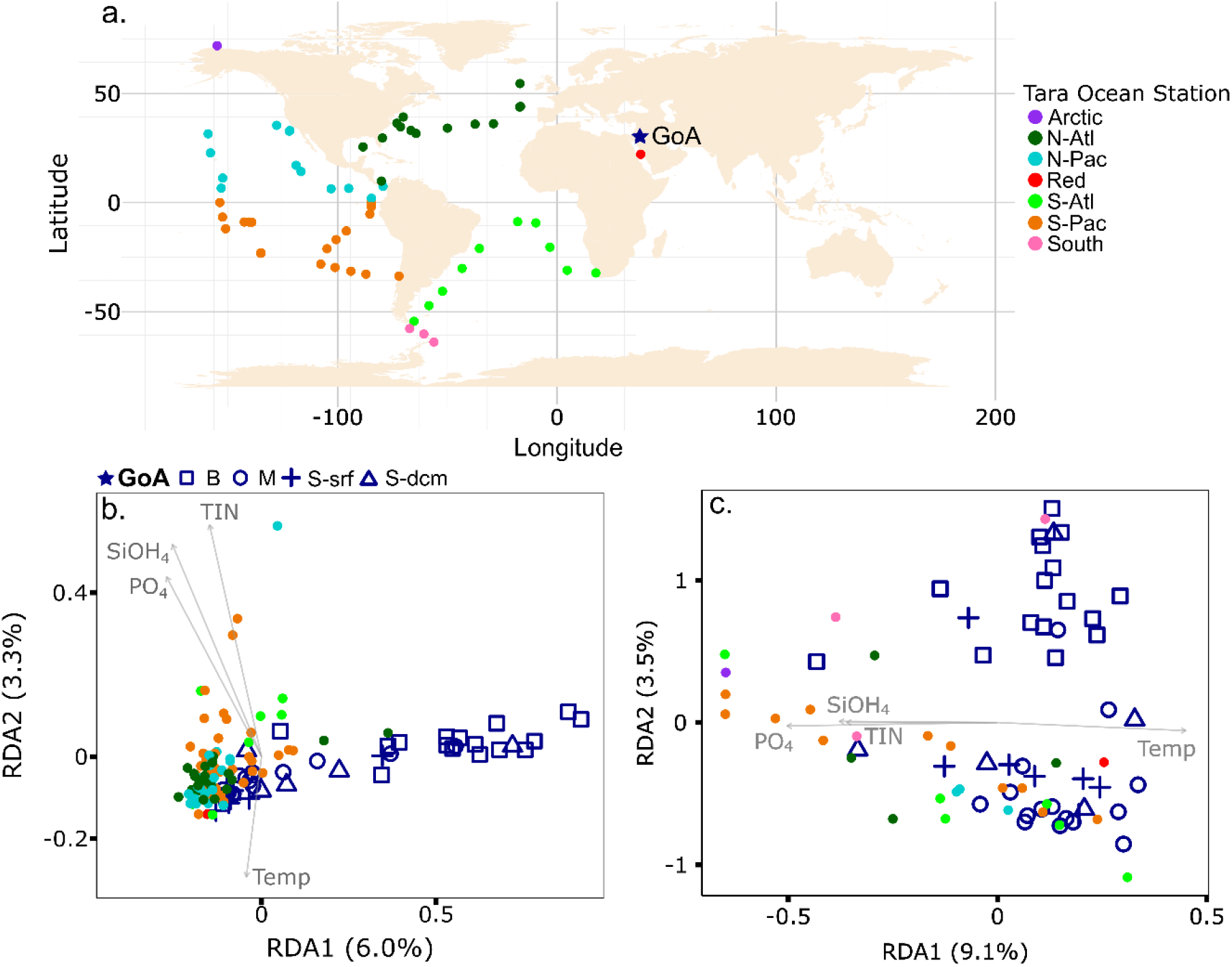
Global-scale community composition of the GoA and *Tara* Oceans samples. (a) Geographical location of the GoA and Tara stations that were examined. RDA analysis was based on the average (duplicates) relative abundance of (b) genera (or lower taxonomic resolution when genus was not assigned) and (c) functional groups at each sampling point. Shapes correspond to the GoA’s seasonal period: B=bloom, M=mixing, S-srf=summer-surface, S-dcm=summer-DCM. Filled circles represent different oceanic basins and seas covered by the *Tara* Oceans cruise: Arctic=Arctic Ocean, N-Atl=North Atlantic, N-Pac=North Pacific, Red=Red Sea, S-Atl=South Atlantic, S-Pac=South Pacific, South=Southern Ocean. Grey arrows show influential environmental variables (p. value <0.05): TIN= total inorganic nitrogen, SiOH4= silica, PO4=phosphate, Temp=temperature. Percentage in parenthesis refers to the explained variability of environmental parameters by each axis.

To test the relationship in functional diversity of protists in the GoA relative to other oceanic settings, we selected a variety of open-ocean and oligotrophic regions that showed high taxonomic similarity to the GoA from the *Tara* Oceans dataset, and also included additional representative *Tara* Oceans nutrient-rich stations from the Arctic and Southern Oceans (Fig. 7c). Additionally, we annotated the main genera of these stations based on the trait table as described above and created a new set of six functional groups that encompassed both *Tara* Oceans and the GoA datasets (Supp. Table 2). RDA analysis of the functional group composition revealed that environmental drivers (water temperature and macronutrients) accounted for about 13% of the distance between assemblages (Fig. 7c). Again, samples from the GoA mixing and stratified surface water periods were mostly similar in their functional assemblage to both open-ocean and Southern Ocean stations associated with high water temperatures and low nutrients. Bloom samples were dissimilar to other regions in their functional structure and were not strongly associated with water temperature and macronutrients.

## Discussion

A marked sucession in protist diversity and functional groups was detected in the Gulf of Aqaba, which occurred in-phase with the local seasonal alternation between oligotrophic and mesotrophic oceanographic conditions. Warmer, nutrient-limited conditions during the stratified spring/summer period favored highly diverse assemblages dominated by protists with heterotrophic functions. These assemblages were established rapidly following nutrient depletion and estabilishment of a stratified water-column during May, and included pico-parasites affiliated mostly with the Syndiniales. This is a hyperdiverse lineage of protistan endoparasites comprising five main taxonomic groups (Groups I-V; Guillou et al., 2008; Rizos et al., 2023) which infect host cells as a minute motile spore (dinospore), multiply intracellularly, and finally kill the host cell before release (Coats & Park, 2002; Chambouvet et al., 2008; Decelle et al., 2022). Although all Syndiniales groups were detected in the GoA, groups I and II were the most prevalent. Similar patterns have been reported from other marine locations (e.g., de Vargas et al., 2015; Ollison et al., 2021; James et al., 2022; Nagarkar & Palenik, 2023; Rizos et al., 2023; Anderson et al., 2024), where Syndiniales groups I and II can infect a wide spectrum of hosts, including other protists (dinoflagellates, cercozoans, radiolarians) and metazoans (copepods, fish eggs) (Anderson et al., 2024). High host specificity has been described in Syndiniales group II, which has been shown to contribute to the regulation of dinoflagellate blooms in nutrient-rich, coastal areas (Chambouvet et al., 2008). In the GoA high Syndiniales abundance and diversity were not concomitant to algal blooms, but occurred instead during the low cell concentrations periods of the stratified and early mixing periods, likely correlating with the presence of specific hosts. These may include other dinoflagellates that were more frequent during the same period, such as *Gymnodinium, Prorocentrum* and *Tripos*, all of which are infected by Syndiniales (Siano et al., 2011; Christaki et al., 2023; Nagarkar & Palenik, 2023; Anderson et al., 2024). In addition most of the dinoflagellates belonged to the PR2 category *Dinophyceae_XXX* which remains largely undefined (Guillou et al., 2013). Many of these may also serve as hosts for Syndinales (Nagarkar & Palenik, 2023), further explaining the high parasitic prevalence during summer. The relative abundance of parasitic Syndiniales is likely inflated due to the high number of rDNA copies in their genomes (e.g., Santoferrara, 2019; Anderson et al., 2024). However, variations in their density over the annual cycle most likely reflect oscillation in parasitic standing-stocks and their importance during the oligotrophic period in the GoA.

In addition to pico-parasites, we detected an important component of protists with potential mixotrophic abilities (phagotrophic) during the stratified season. These include several dinoflagellate genera such as *Gonyaulax, Torodinium*, and *Tripos (Smalley et al*., *2003; Jeong et al*., *2005; G*ómez, 2009). Such a combination of carbon fixation by photosynthesis with heterotrophy can compensate for the nutrient limitation occuring during summer in the GoA (Stoecker et al., 2017). Furthermore, a contingent of phagotrophic functional-groups (pico- and micro-phagotrophs), corresponding to Marine Stramenopiles (MAST), as well as Oligotrichs and Strombidiidae ciliates, were detected during the same period. These findings agree with observations from other oligotrophic ecosystems (Hartmann et al., 2012), illustrating the prevalence of various heterotrophic modes in stratified oceanic regimes (Mitra et al., 2014; Blanco-Bercial et al., 2022).

The onset of the mixing layer during the fall (around November) triggered a gradual transition in protist communities along with the deepening of the MLD. Over this period, the prevalence of parasitic and potential mixotrophic clades progressively declined, likely following the decline of host availability (although diverse assortments of Syndinales remained, indicating effective parasitic propagation) and increased inorganic-nutrient availability. Reciprocally, the prevalence of small-size (pico and nano) photoautotrophs increased, such that during the bloom they accounted for over 50% of the protist 18S amplicon sequences. These included *Ostreococcus*, a genus from the Chlorophyte Order Mamiellales. This is consistent with previous reports from open-ocean oligotrophic systems, indicating an overall preference of *Ostreococcus* for nutrient-richer conditions, and an association with increased phytoplankton productivity (Treusch et al., 2012; Choi et al., 2020; Eckmann et al., 2024). Other Mamiellales genera (*Micromonas* and *Bathycoccus*) were relatively more important during the stratified period, with *Ostreococcus* still dominating in deeper and nutrient rich DCM layers. This result highlights a consistent differentiation between the main Mamiellales genera and the potential role of phytoplankton standing-stocks inhabiting DCM as a reservoir for winter assemblages in the mixed water column (Eckmann et al., 2024).

Other important phytoplankton clades detected in the GoA and included both in the functional groups of the pico/nano-autotrophs and silicified-autotrophs were the Ochrophyta and Haptophyta, together representing an average of 13% of amplicon sequences across seasons. The contribution of both Phyla was more important during the bloom period. However, clear distinctions were detected between the relatively shallow bloom of 2021 and the larger 2022 bloom following a colder winter and much deeper MLD. During the 2021 bloom, Ochrophyta from the genus *Aureococcus* and pennate diatoms, associated respectively with the pico/nano-autotrophs and the silicified-autotrophs functional groups, dominated. By contrast, during the nutrient-richer bloom of 2022 following deeper mixing, a much more important and diverse contingent of diatoms was observed. These comprised notably taxa from the centric-diatom *Skeletonema* and polar-centric-Mediophyceae, as well as the Haptophyta *Gephyrocapsa huxleyi*, similar to what is frequently detected in nutrient-rich, high latitudes regions (Tréguer et al., 2018; Cascella et al., 2020; Choi et al., 2020), emphasizing the central regulatory role of bottom-up factors in bloom development and community composition. However, we note that the 2022-bloom corresponded to the protist diversity-minimum, suggesting additional underlying competitive exclusion mechanisms influencing the composition of protist communities during bloom development.

The rise of diatoms and coccolithophores in response to nutrient input in the water column can also have larger ecological consequences. The growth of centric diatom species, including some chain-forming taxa, and heavily armored coccolithophores, can boost carbon export rates to the deep ocean during bloom development and senescence (Tréguer et al., 2018; Li et al., 2024). Consistent with this, carbon export rates in the GoA increase as a function of mixing depth and bloom size (Torfstein et al., 2020; Keuter et al., 2023). Interestingly, the armored-micro-phagotrophs increased also during the 2022 bloom. This functional group is composed of Radiolaria, an ancient eukaryotic lineage of heterotrophic and photosymbiotic mixotrophic taxa (Decelle et al., 2015), whose production has been shown to correlate with primary productivity and can markedly influence remineralization and carbon export rates in the oceans (Takahashi, 1987; Gutierrez-Rodriguez et al., 2019; Qu et al., 2022; Laget et al., 2024). Radiolaria abundance peaks in the GoA may thus represent period of higher productivity and export fluxes.

Similarities between the GoA and other oligotrophic regions of the world ocean are often evoked in the literature (Reiss & Hottinger, 1984; Lindell & Post, 1995; Labiosa et al., 2003). If this may be the case from an oceanographic perspective, the relation of protist assemblages inhabiting the GoA to those from other oceanic regions remains unclear. To see whether, from a biological viewpoint the GoA represents a typical oligotrophic ecosystem, we compared our dataset to similar amplicon data collected worldwide during the *Tara* Oceans circum-global expedition (de Vargas et al., 2015). Protist assemblages (from the pico-nano plankton) inhabiting surface water during the stratified period of the GoA indeed resemble those of other open ocean ecosystems, with Syndiniales and Dinophyceae as the main groups (de Vargas et al., 2015; Pierella Karlusich et al., 2020). In contrast, assemblages observed during the bloom and to some extent at the DCM were distinct from other oceanic samples. This could result from the development of singular subtropical communities in response to unique oceanographic conditions in the GoA. However, this result may also suffer from the low representation of coastal regions and limited temporal scope within the *Tara* Oceans data. Further comparisons with coastal, upwelling, and high-nutrient high-latitude regions is required for a better assessment of the relation of winter protists communities in the GoA to other global regions.

## Conclusions and Perspectives

We have provided in this study the first detailed synthesis of the functional diversity and seasonality of protist assemblages from the pico-nano plankton in the epipelagic sunlit layer of the subtropical Gulf of Aqaba. A pronounced seasonality in the taxonomic and functional attributes of protist communities was detected. In summary, we observed a major transition between communities largely dominated by heterotrophs, notably parasites, during the oligotrophic periods, and communities dominated by phototrophs during winter and the Spring bloom periods when nutrient availability increased. Moreover, a strong interannual variation in bloom formation was measured, linked to differential mixing depths and consequent nutrient loads available to primary prodcuers. Therefore, the GoA planktonic ecosystem is marked by seasonal and interannual shifts in trophic ecology and energy fluxes, which likely influence local ecology and biogeochemistry over the annual cycle (Goericke, 1998; Lomas et al., 2013). Furthermore, close similarities between planktonic assemblages from the GoA and other (sub)tropical regions from the world’s open ocean were detected, indicating that the GoA, when stratified during the summer, can serve as a model for the study of oligotrophic ecosystems. As such our study adds to the current effort to better understand the ecology of oligotrophic marine ecosystems. However, the ecological succession presented here across oligotrophic-mesotrophic gradients in the GoA can also represent a case study simulating the responses of high-latitude communities to ongoing oligotrophication due to the expansion of stratified regions under global warming. Future coupled quantification of the contribution of specific groups to primary production and export fluxes, as well as their functional role, will enable better mechanistic understanding and modelling of ecological processes and dynamics in subtropical current and future ecosystems.

## Acknowledgments

This study was supported by the Israel Science Foundation grant 2921/20, attributed to MJF and by a PhD fellowship from the Interuniversity Institute for Marine Sciences in Eilat attributed to YA. MR-B was funded by the Israel Ministry of Science and Technology grant 001126. We would like to thank Tanya Rivlin for nutrient measurements, the crew from the R/V Sam Rothberg for assistance during sampling as well as Sarah Romac for sharing DNA extraction protocols.

## Data Availability Statement

The raw 18S rRNA amplicon data for this study can befound in the National Centre for Biotechnology Information (NCBI) database under BioProject PRJNA1169359: https://www.ncbi.nlm.nih.gov/bioproject/?term=PRJNA1169359

## Author contributions

**Yoav Avrahami:** conceptualization; data curation; formal analysis; investigation; methodology; software; validation; visualization; writing – original draft; writing – review & editing. **Raffaele Siano:** methodology; validation; writing – review & editing. **Max Rubin-Blum:** methodology; validation; writing – review & editing. **Gil Koplovitz:** investigation; project administration. **Nicolas Henry:** methodology; validation. **Colomban de Vargas:** resources; writing – review & editing. **Miguel J. Frada:** conceptualization; formal analysis; funding acquisition; project administration; resources; supervision; validation; writing – original draft; writing – review & editing.

