## Supplementary Figures 1-4 for "Switch in parasitic and autotrophic-dominated protist assemblages coupled to seasonal oligotrophic-mesotrophic gradients in the sunlit layer of a subtropical marine ecosystem"

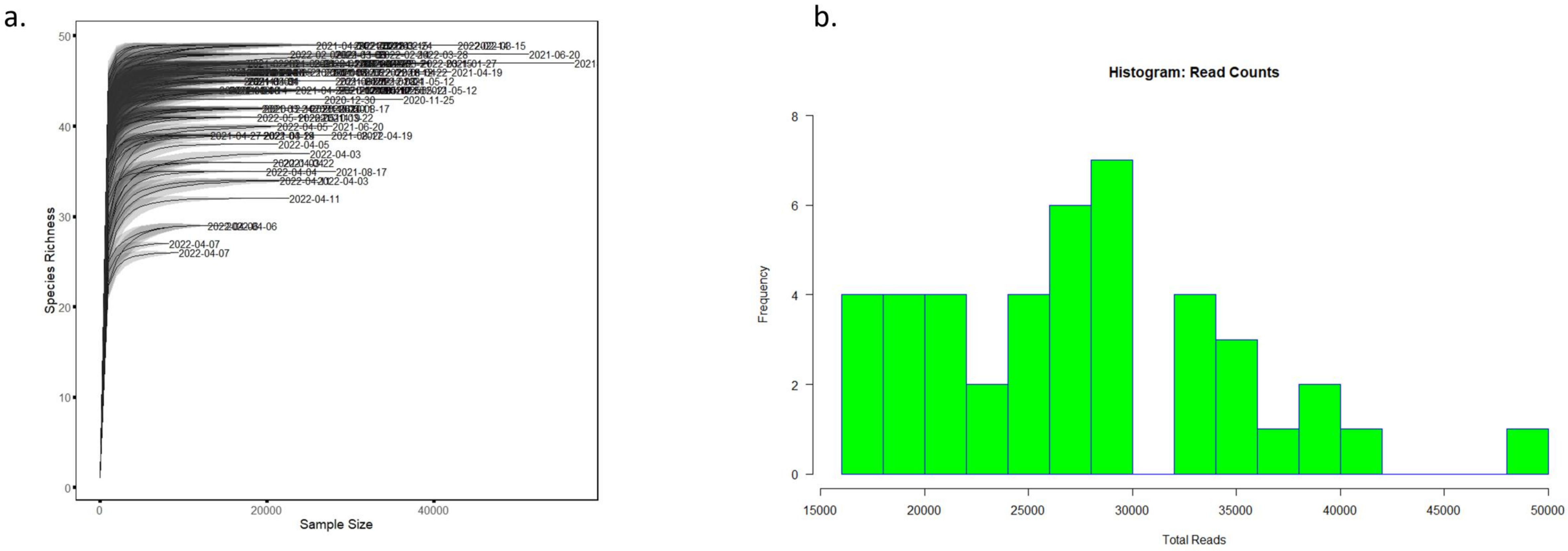


**Supplementary Figure 1.** ASV coverage of all sampling points. (a) Rarefaction curve of ASV richness. Labels denote the date of samping (yyyy-mm-dd). (b) Histogram of total ASV reads in each of the sampling points.

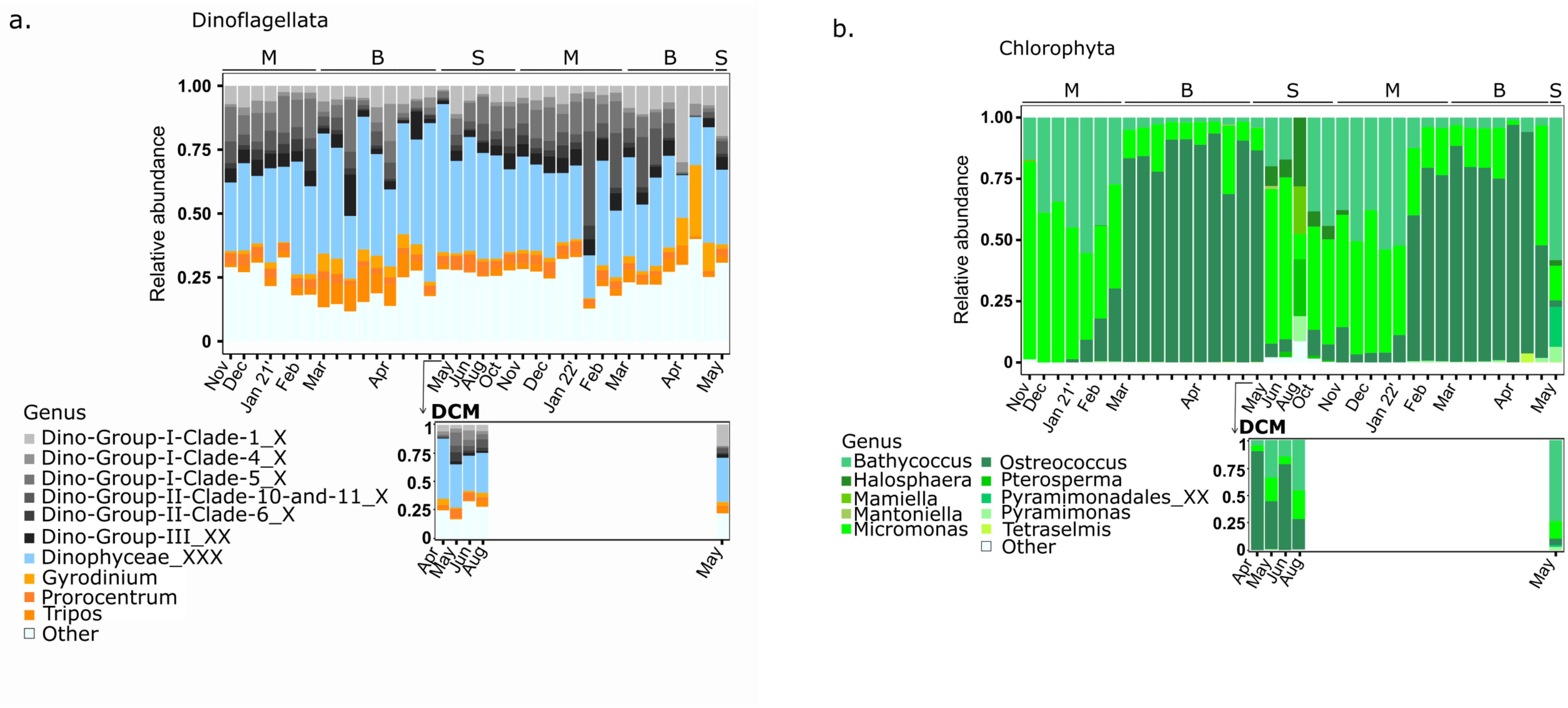


**Supplementary Figure 2.** Main genera of the phyla Dinoflagellata (a) and Chlorophyta (b) in the Gulf of Aqaba. Relative abundance of the top 10 most abundant genera in surface waters (upper panel) and the corresponding main genera at the DCM during summer (lower panel), based on ASV's average reads (duplicates). Letters in the top panel represent seasonal periods: M=mixing, B=bloom, S=summer stratification.

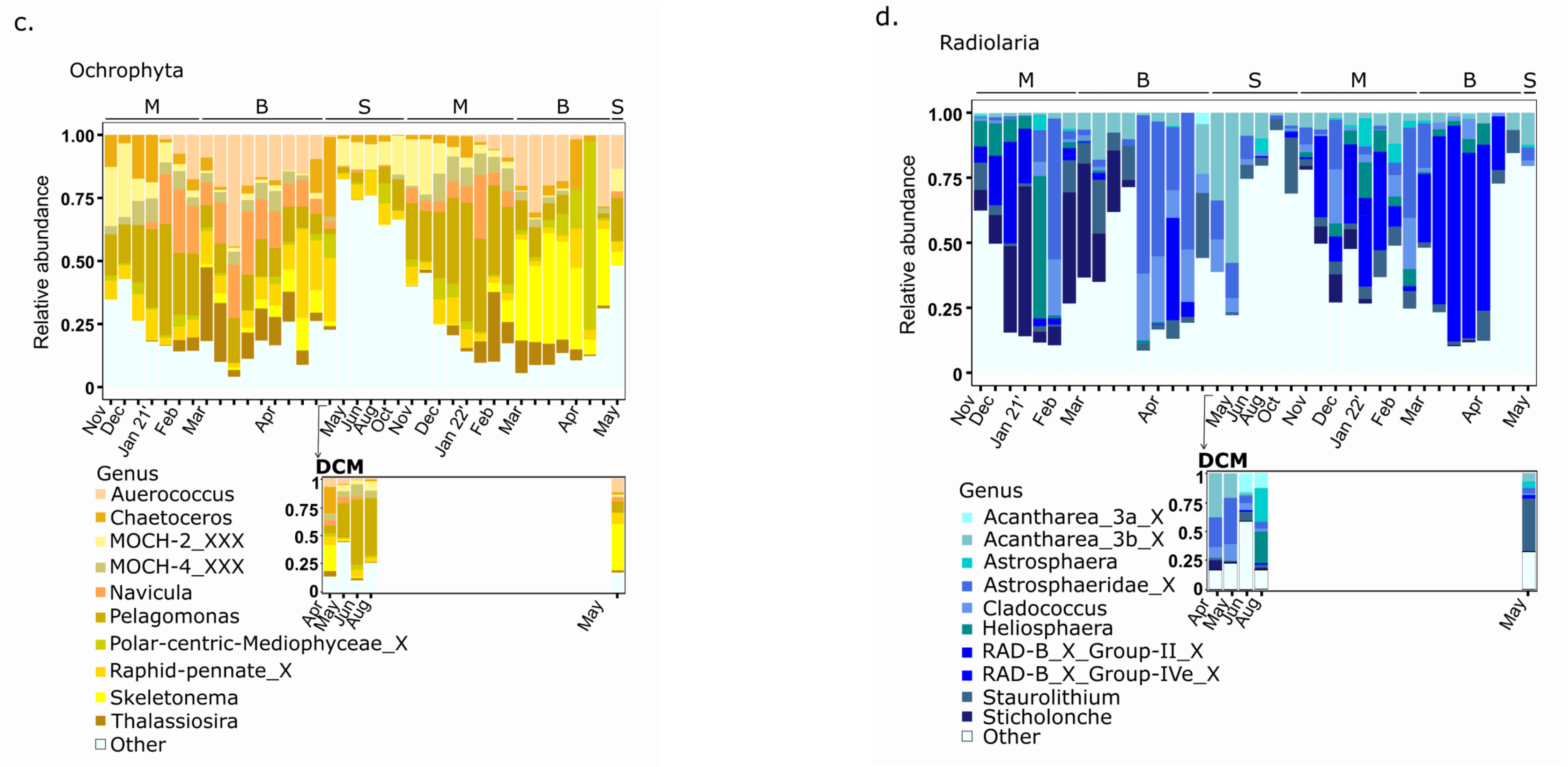


**Supplementary Figure 2.** Main genera of the phyla Ochrophyta (c) and Radiolaria (d) in the Gulf of Aqaba. Relative abundance of the top 10 most abundant genera in surface waters (upper panel) and the corresponding main genera at the DCM during summer (lower panel), based on ASV's average reads (duplicates). Letters in the top panel represent seasonal periods: M=mixing, B=bloom, S=summer stratification.

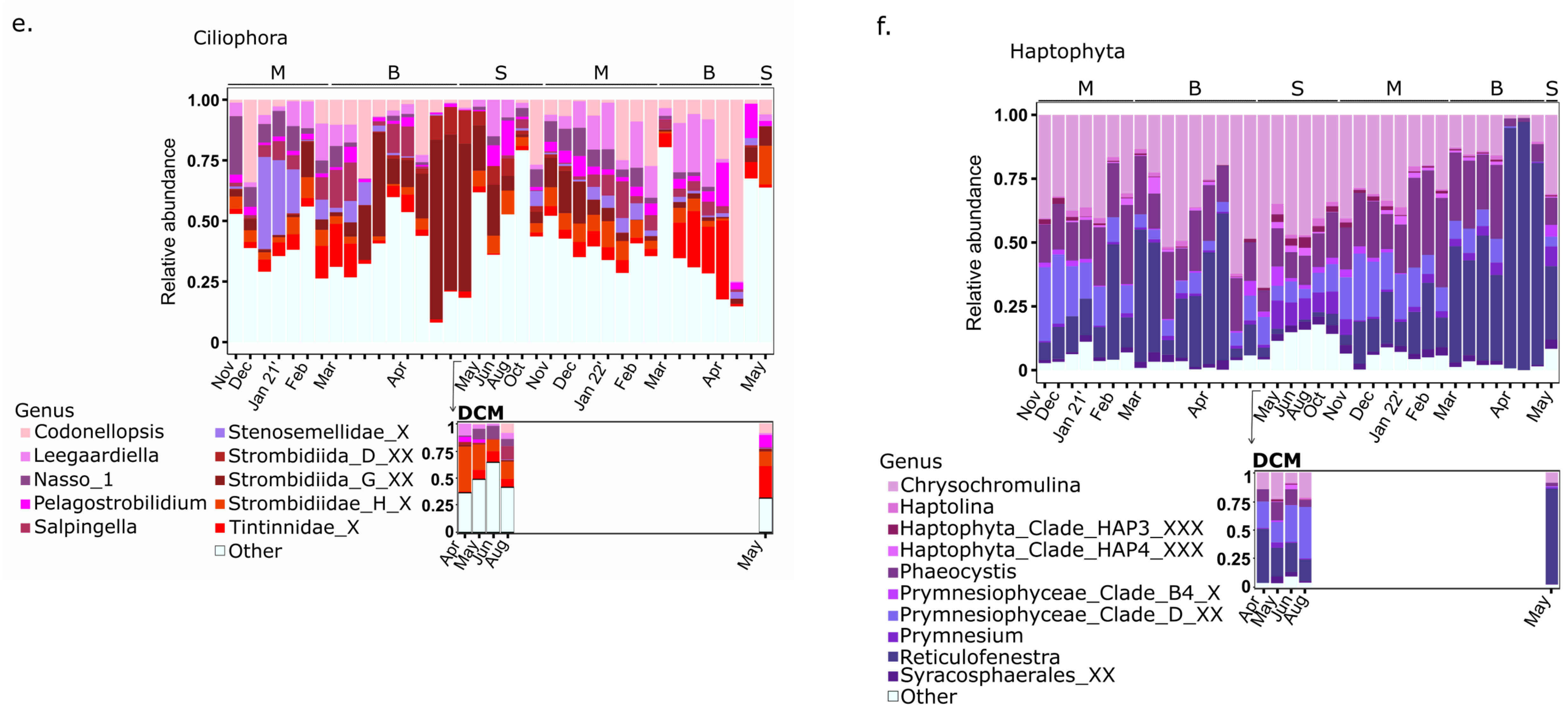


**Supplementary Figure 2.** Main genera of the phyla Ciliophora (e) and Haptophyta (f) in the Gulf of Aqaba. Relative abundance of the top 10 most abundant genera in surface waters (upper panel) and the corresponding main genera at the DCM during summer (lower panel), based on ASV's average reads (duplicates). Letters in the top panel represent seasonal periods: M=mixing, B=bloom, S=summer stratification.

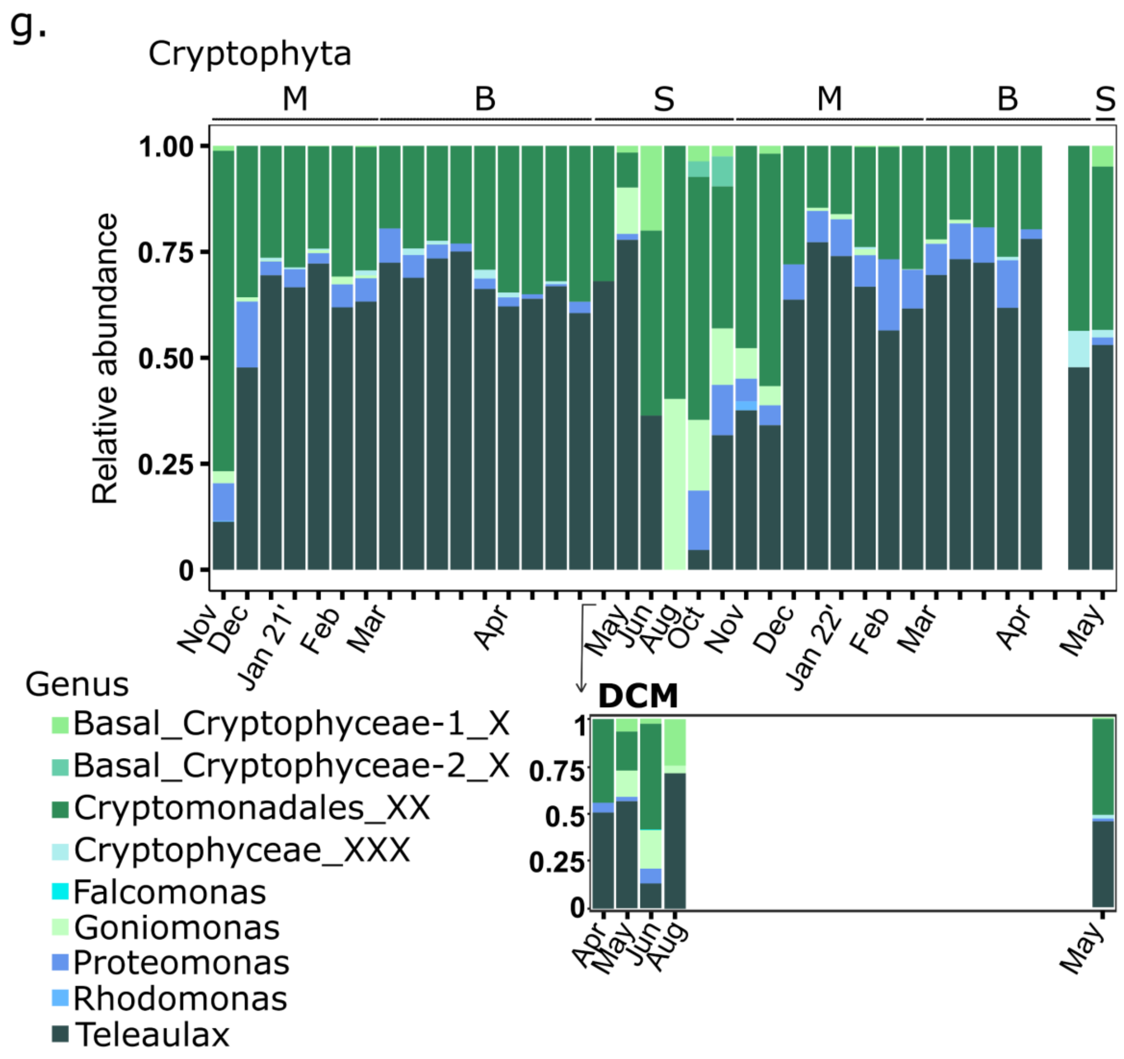


**Supplementary Figure 2.** Main genera of the phyla Cryptophyta (g) in the Gulf of Aqaba. Relative abundance of the top 10 most abundant genera in surface waters (upper panel) and the corresponding main genera at the DCM during summer (lower panel), based on ASV's average reads (duplicates). Letters in the top panel represent seasonal periods: M=mixing, B=bloom, S=summer stratification.

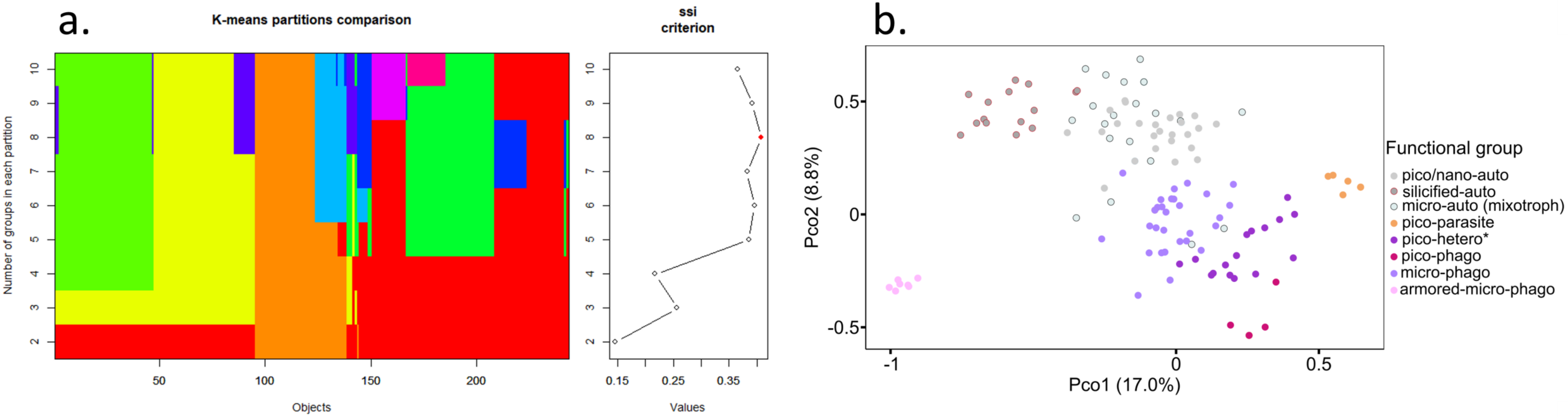


**Supplementary Figure 3.** Clustering of functional groups. (a) K-means and ssi criterion suggest assignment of genera to 8 functional groups. (b) PCoA analysis based on Gower distance of the selected 243 genera. Each circle represent a genus, distances are based on trait-composition. Colors show the functional groups. Percentage in parenthesis refers to the explained variability that each axis prvoides.

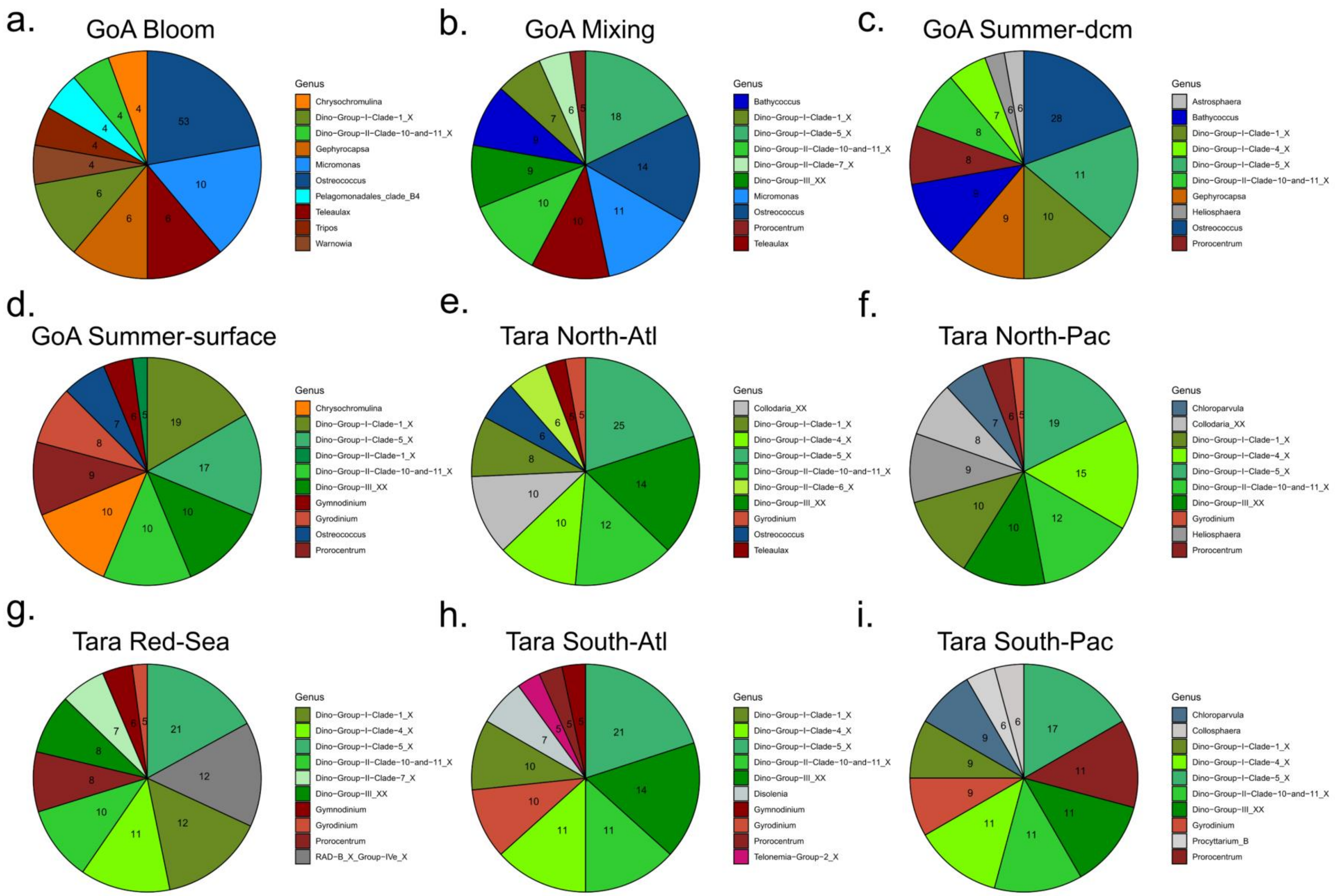


**Supplementary Figure 4.** Pie charts of the top 10 main genera at each sampling station. Numbers represent relative abundance (%, 1-100). (a) GoA bloom. (b) GoA mixing. (c) GoA summer-DCM. (d) GoA summer-surface. (e) Tara North-Atlantic. (f) Tara North-Pacific. (g) Tara Red-Sea. (h) Tara South-Atlantic. (i) Tara South-Pacific
